# Alpha/Beta Hydrolase Domain-Containing Protein 2 regulates the rhythm of follicular maturation and estrous stages of the female reproductive cycle

**DOI:** 10.1101/684951

**Authors:** Ida Björkgren, Dong Hwa Chung, Sarah Mendoza, Liliya Gabelev-Khasin, Andrew Modzelewski, Lin He, Polina V. Lishko

## Abstract

Therian female fertility is defined by a successful and strictly periodic ovarian cycle, which is under the control of gonadotropins and steroid hormones, particularly progesterone and estrogen. The latter two are produced by the ovaries that are engaged in controlled follicular growth, maturation and release of the eggs, i.e. ovulation. It is well known that steroid hormones regulate ovarian cycles via genomic signaling, by altering gene transcription and protein synthesis. However, despite this well-studied mechanism, steroid hormones can also signal via direct, non-genomic action, by binding to their membrane receptors. Here we show, that the recently discovered sperm membrane progesterone receptor α/β hydrolase domain-containing protein 2 (ABHD2) is highly expressed in mammalian ovaries where the protein plays a novel regulatory role in follicle maturation and the sexual cycle of females. Ablation of *Abhd2* caused a dysregulation of the estrous cycle rhythm with females showing shortened luteal stages while remaining in the estrus stage for a longer time. Interestingly, the ovaries of *Abhd2* knockout (KO) females resemble polycystic ovary morphology with a high number of atretic antral follicles that could be rescued with injection of gonadotropins. Such a procedure also allowed *Abhd2* KO females to ovulate a significantly increased number of mature and fertile eggs in comparison to their wild-type littermates. These results suggest a novel regulatory role of ABHD2 as an important factor in non-genomic steroid regulation of the female reproductive cycle.

## Introduction

Female fertility is highly regulated by the hypothalamic-pituitary-gonadal axis of hormone secretion, which results in a balanced estrogen to progesterone signaling and, in turn, produces the cyclic nature of ovarian follicle development. In many placental mammals this cycle is known as the estrous cycle, with the exception of primates which have a menstrual cycle. The estrous cycle is generally subdivided into four phases: proestrus, estrus, metestrus and diestrus (Supplemental Figure S1). The first phase - proestrus- begins with the rapid growth of several ovarian follicles and can, in mice, last about one day. During this phase the old corpus luteum (CL) degenerates, while the vaginal epithelium proliferates. Proestrus can be easily identified by the large number of non-cornified nucleated epithelial cells found in smears collected by vaginal lavage (Supplemental Figure S1). Steadily increasing levels of estrogen stimulate preovulatory follicles that undergo their final growth phase. The second phase, estrus, follows proestrus and is the stage that experiences the peak of gonadotropin secretion by the pituitary gland, resulting in maximal secretion of estrogens by the ovaries. Dominant follicles produce high levels of estradiol to inhibit the development of smaller antral follicles^1^ and simultaneously promote ovulation through expression of the nuclear progesterone receptor and induction of progesterone secretion^2,3^. Ovulation occurs during the estrus stage, which together with proestrus comprise the follicular phase. The estrus stage can also be easily identified using vaginal smears that show presence of larger cornified epithelial cells (Supplemental Figure S1). Progesterone and its receptors are primarily needed during the subsequent luteal phase (metestrus and diestrus), which occurs immediately after ovulation, and results in the consequent development of CL from the ovulated follicle (Supplemental Figure S1). CL is the main producer of progesterone, which peaks during diestrus^4^, and if pregnancy is not obtained, the CL goes through luteolysis, which leads to reduced progesterone secretion and stimulates development of the immature follicles.

Progesterone is a powerful regulator of the ovarian cycle that occurs via genomic signaling mechanisms and results from progesterone binding to its nuclear receptors inside the cell, and leads to changes in gene transcription and protein synthesis. However, it has been known that progesterone may also signal via a non-genomic or direct pathway, by binding to its membrane receptors. The latter signaling pathway is required for frog oocyte maturation^5^, human sperm cell activation^6–8^, and likely triggers anesthesia in rodents^9^. Recently, by using transcriptionally silent spermatozoa as a model, we have identified the novel membrane progesterone receptor, the α/β hydrolase domain–containing protein 2 (ABHD2) as well as described the signaling pathway that is initiated by progesterone association with ABHD2^10^. Monoacylglycerol lipase ABHD2 is the first evolutionary conserved steroid-activating enzyme that hydrolyzes endocannabinoid 2-arachidonoylglycerol^10^. Here, we show that ABHD2 is not only expressed in sperm, but also display high expression in ovaries, particularly in CL and stromal cells. To study this further, we have generated an ABHD2 knockout mouse by using the highly efficient CRISPR Ribonucleoprotein Electroporation of Zygotes (CRISPR-EZ) technique^11,12^ and evaluated its fertility phenotype. We show that ABHD2 is needed to regulate the cyclic maturation of follicles. The reproductive cycle of *Abhd2^−/−^* females shows a prolonged estrus stage while the time spent in luteal phase and proestrus is shortened. Follicle development was also affected with a higher number of follicles reaching antral stage. However, a majority of the follicles showed signs of atresia and the *Abhd2^−/−^* females produced similar numbers of ovulated eggs and litter sizes as wild-type females. Interestingly, when *Abhd2* KO females were injected with hormones to induce superovulation, the atretic antral follicles were rescued and the females ovulated a significantly higher number of mature and fertile eggs in comparison to their wild-type littermates. This phenotype is similar to that of polycystic ovarian syndrome where atretic follicles can be induced to ovulate through hormone injections. However, the increased number of follicles reaching antral stage did not cause a depletion of the ovarian reserve, as 8-month-old mice, heterozygous or homozygous for *Abhd2* ablation, were still able to produce similar number of eggs as wild-type females. The results indicate that this evolutionary conserved protein is an important regulator of the female reproductive cycle.

## Results

### *Abhd2* expression in ovarian stromal cells and corpora lutea

While ABHD2 has been described as an important regulator of sperm function^10^, the function of ABHD2 in female reproduction was not known. Interestingly, unlike the male reproductive tissues, we found that ABHD2 is not detected in the female gametes but is predominantly expressed in the stromal cells surrounding the developing follicles (Fig. 1a). The presence of both ABHD2 protein and mRNA was observed in pre-pubertal mouse ovaries, as shown here by immunohistochemical (IHC) staining and reverse transcriptase quantitative PCR (RT-qPCR; Fig. 1b and c). Around one month after birth, when the ovulatory cycle begins, ABHD2 expression is further found in the lutein cells of corpus luteum (Fig. 1a). In correlation with this, qPCR studies showed highest *Abhd2* mRNA presence right after ovulation during the estrus stage (Fig. 1c), and the expression levels are even further increased after induction of superovulation by gonadotropin injection (Fig. 1c).

**Fig. 1.**
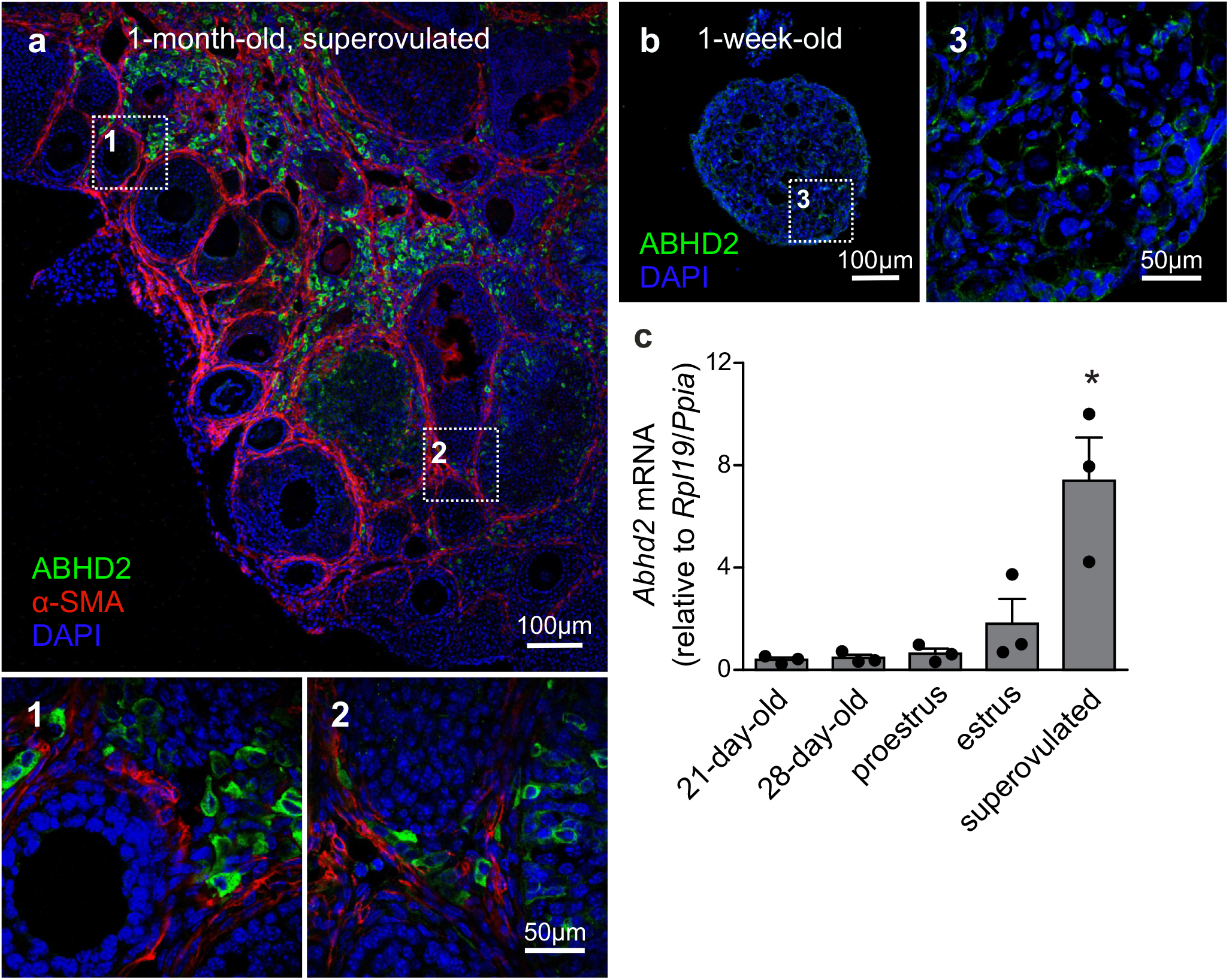
Expression of Abhd2 in the mouse ovary. **a)** Immunohistochemical staining of a one-month-old superovulated wild-type mouse ovary shows ABHD2 in the stromal cells (1) and in cells of the corpus luteum (2) while the protein is not present in the granulosa cells and oocytes of developing follicles, demarcated by staining of the surrounding smooth muscle layer. **b)** and 3) Stromal cells of 1-week-old pre-pubertal mouse ovaries also display ABHD2 staining while **(c)** shows similar expression levels of *Abhd2* before and after the first round of ovulation, as detected in 21- and 28-day-old mice ovaries. The expression levels increase after ovulation, as seen in ovaries at estrus and after superovulation. Samples (n=3 for each timepoint) were collected from 3-month-old females in proestrus and estrus and 1-month-old superovulated females. Expression levels measured by qPCR were normalized to the expression of ribosomal protein L19 (*Rpl19*) and peptidylprolyl isomerase A (*Ppia*). Statistical significance was calculated using the unpaired t-test and the significance of changes is indicated as follows: *, *p ≤ 0.05*.

### Generation of the *Abhd2* knockout mouse line

To study the role of ABHD2 in female reproduction, we have generated an *Abhd2* knockout mouse line by utilizing the CRISPR-EZ technique^11^, where single guide RNA (sgRNA)/Cas9 complexes are delivered into mouse zygotes by electroporation. Cas9-mediated deletion of *Abhd2* exon 6 was performed through intron specific binding of two sgRNAs (Fig. 2a). The deletion resulted in the removal of a serine (S208), the amino acid important for the catalytic function of ABHD2, and resulted in a frameshift and a premature stop codon in exon 7 (Supplemental Fig. S2). Complete ablation of *Abhd2* was further confirmed by genotyping PCR (Fig. 2b), qPCR (Fig. 2c), western blotting (Fig. 2d) and IHC analysis of the mouse ovary (Fig. 2e-f). The knockout mice were born at a Mendelian ratio when breeding heterozygous males with females. No obvious morphological or health phenotypes have been observed, and both homozygous and heterozygous females appeared fertile and healthy with similar body weight as their wild-type littermates (Supplemental Fig. S3a).

**Fig. 2.**
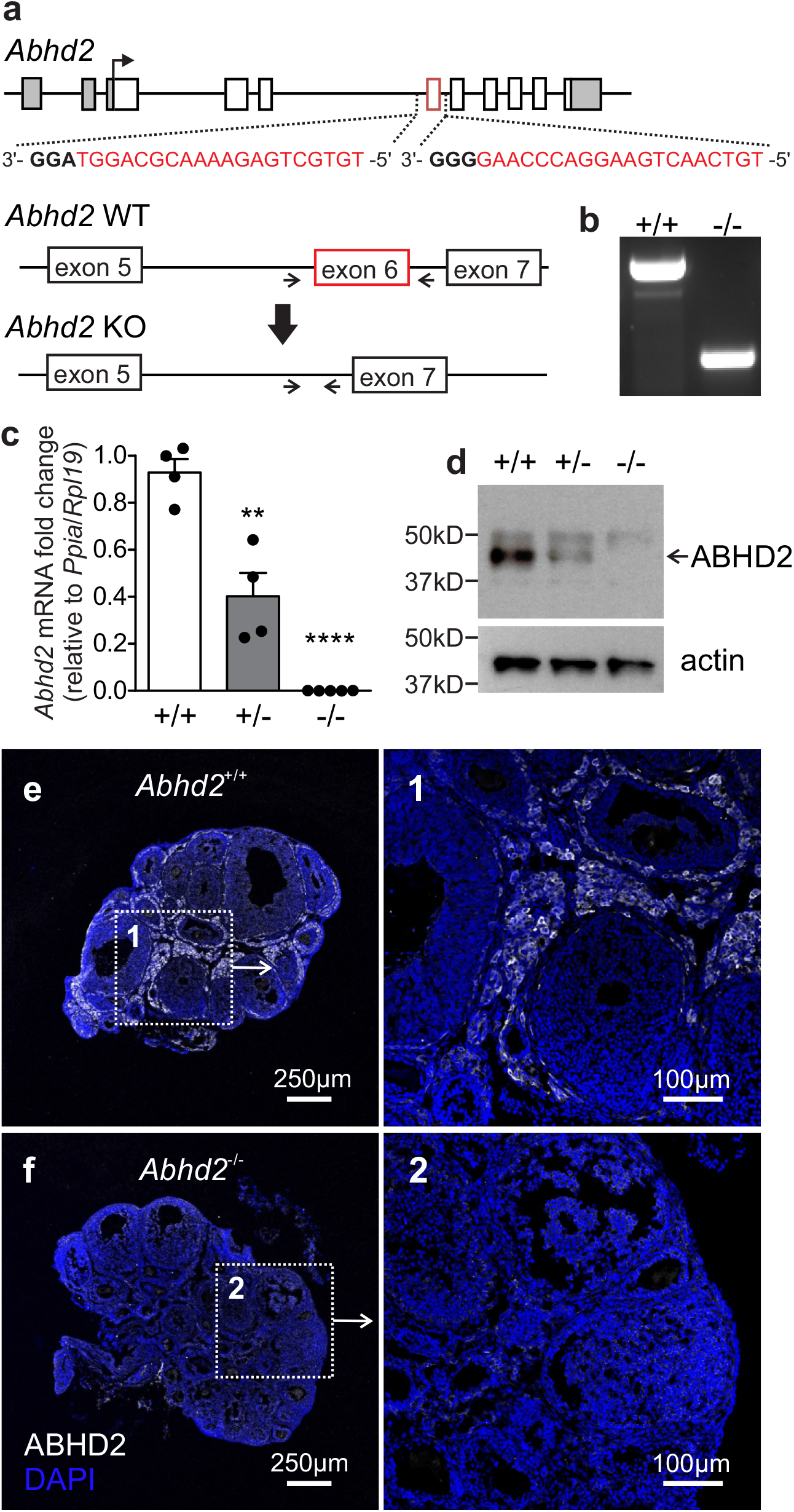
Generation of the Abhd2 knockout mouse line. **a)** Schematic picture of the *Mus musculus Abhd2* gene with untranslated regions marked in grey. Deletion of exon 6 (in red) was performed by CRISPR/Cas9, targeting sequences in the surrounding introns. The sequences of the two sgRNAs are labeled in red and the PAM sequence in bold. **b)** Genotyping of mice was performed using primers that bind outside the deleted sequence (arrows). The PCR product from a wild-type mouse (+/+) contains the exon 6 sequence while the full knockout (−/−) lacks *Abhd2* exon 6. *Abhd2^+/−^* mice show a significant reduction of *Abhd2* expression in both **c)** qPCR and **d)** western blot studies while the full knockout *Abhd2^−/−^* completely lacks expression of exon 6 and the protein. Expression levels detected by qPCR were normalized to the expression of ribosomal protein L19 (*Rpl19*) and peptidylprolyl isomerase A (*Ppia*). Statistical significance was calculated using the unpaired t-test and the significance of changes is indicated as follows: **,*p ≤ 0.01;* ****,*p ≤ 0.0001*. **e)** The *Abhd2^+/+^* mouse ovary shows ABHD2 in the stromal cells surrounding the follicles. F) No protein staining was detected in the *Abhd2^−/−^* ovary.

### Superovulation of *Abhd2^+/−^* and *Abhd2^−/−^* females gives rise to an increased number of ovulated eggs

Both the litter size and birth rate of *Abhd2^+/−^* and *Abhd2^−/−^* females were not significantly different from that of wild-type mice (Table 1). In addition, spontaneous ovulation gave rise to similar numbers of eggs for all genotypes (Fig. 3a). However, when injected with gonadotropins: pregnant mare serum gonadotropin (PMSG) and human chorionic gonadotropin (hCG)- a standard method to induce superovulation^13^, both the *Abhd2^+/−^* and *Abhd2^−/−^* females produced significantly higher numbers of ovulated eggs (Fig. 3b). The phenotype was not a result of prematurely ovulated follicles, as *in vitro* fertilization studies led to a similar success rate in formation of blastulae as when using eggs from wildtype females (*Abhd2^+/+^*: 72.4% ± 11.3% n=3, *Abhd2^+/−^*: 81.5% ± 9.1%, *Abhd2^−/−^*: 75.0% ± 8.3% fertilized eggs, n=4 for both genotypes). The fertility of 7-8-month-old *Abhd2^+/−^* and *Abhd2^−/−^* females in normal breeding was similar to that of wild-type mice (Table 1). Superovulation of these older animals also gave rise to similar numbers of ovulated eggs although some heterozygous mice still showed numbers comparable to that of much younger animals (Fig. 3c).

**Fig. 3.**
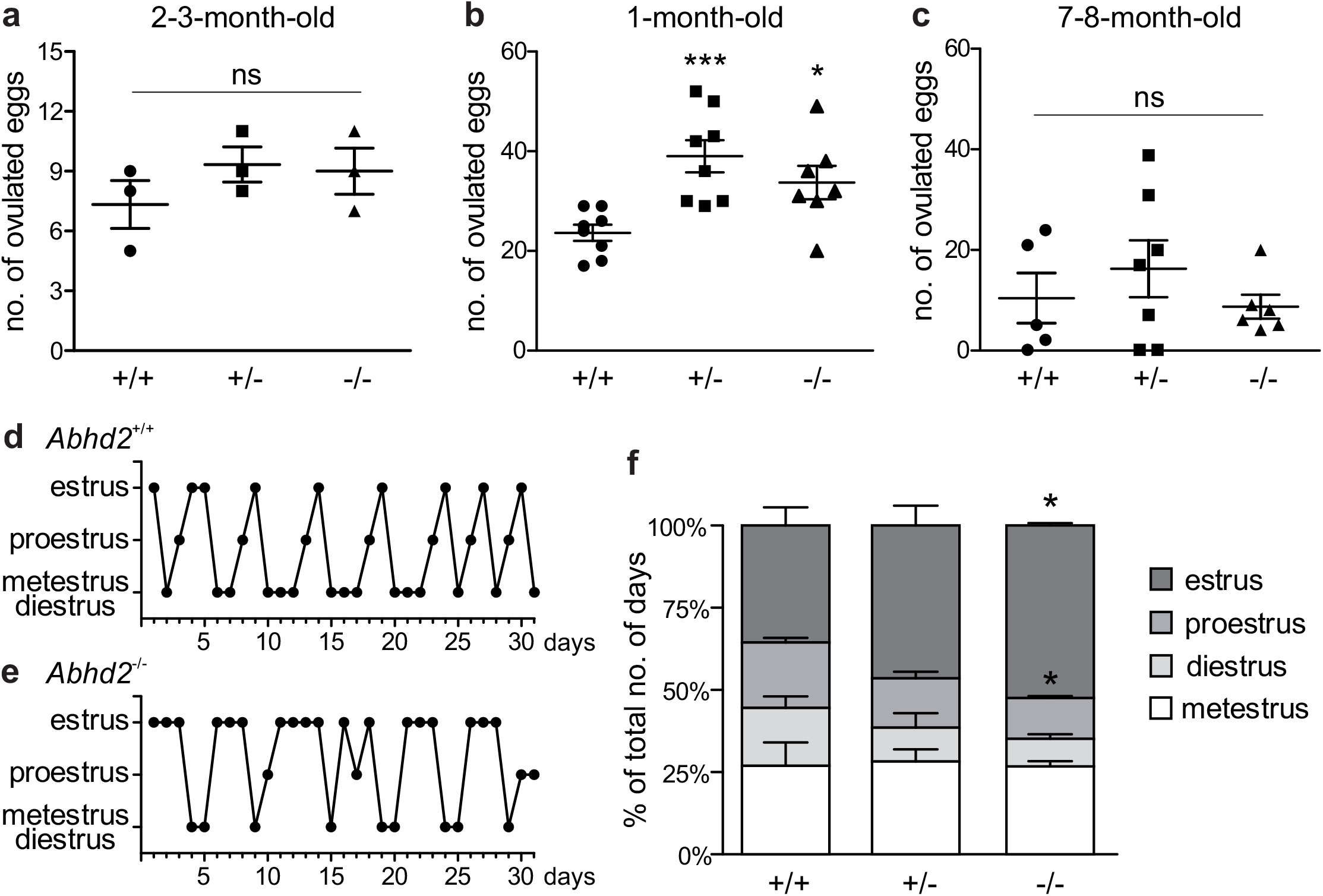
Abhd2^−/−^ female ovulation and estrous cycle. **a)** Spontaneous ovulation gave rise to similar numbers of eggs for all genotypes. **b)** *Abhd2* ablation caused significantly increased numbers of ovulated eggs after induction of superovulation, in both 1-month-old *Abhd2^+/−^* and *Abhd2^−/−^* females compared to *Abhd2^+/+^* mice. **c)** Superovulation of 7-8-month-old females gave rise to similar numbers of ovulated eggs among genotypes. Statistical significance was calculated using the unpaired t-test and the significance of changes is indicated as follows: *, *p ≤ 0.05*; ***, *p ≤ 0.001*. **d)** Representative graphs of the estrous cycle of a **d)** *Abhd2^+/+^* and **e)** *Abhd2^−/−^* female mouse where the *Abhd2* knockout female shows prolonged stages in estrus although displaying cyclical changes throughout the month. **f)** The combined measurements of three mice of each genotype also showed a significantly altered distribution of days in the different estrous stages, in particular the increased percentage of days in estrus and decreased percentage of days in proestrus of *Abhd2^−/−^* females compared with *Abhd2^+/+^* mice.

**Table 1.** Breeding efficiency of *Abhd2^−/−^* females

|  |  |  |  |  |
| --- | --- | --- | --- | --- |
| Female genotype | +/+ | +/- | -/- | -/- |
| Male genotype | +/+ | +/- | +/+ | +/- |
| No. of breeding pairs | 3 | 9 | 2 | 5 |
| Pups/litter | 5.92 ± 0.72 | 6.46 ± 0.25 | 6.73 ± 0.68 | 5.88 ± 0.57 |
| Days between litters | 32.4 ± 3.27 | 31.6 ± 2.03 | 31.3 ± 3.00 | 26.1 ± 1.91 |

### *Abhd2* ablation results in a dysregulated estrous cycle

Interestingly, when following the estrous cycle of virgin females for one month, the *Abhd2^−/−^* females presented with a significantly higher percentage of days in estrus while the proestrus and diestrus stages were shortened compared to wild-type females (Fig. 3d-f). Although the *Abhd2^−/−^* mice showed a prolonged estrus stage they cycled in a regular fashion (*Abhd2^+/+^*: 4.75 ± 0.33 days/cycle, *Abhd2^+/−^*: 4.8 ± 0.53 days/cycle, *Abhd2^−/−^*: 4.43 ± 0.29 days/cycle) and gave rise to litters at a similar rate as *Abhd2^+/+^* females (Table 1). Previous studies have shown that altered estrogen and androgen levels during pregnancy can affect the estrous cycle of female pups^14^. To determine whether or not the maternal genotype influenced the observed phenotype, the estrous cycle of *Abhd2^+/−^* females born from wild-type mothers who were mated with homozygous males or homozygous mothers mated with heterozygous males were compared. The month-long measurements showed a similar prolonged estrus stage in all females (*Abhd2^+/−^* females born from *Abhd2^+/+^* mothers: 47.2% ± 3.7%; *Abhd2^+/−^* mothers: 46.5% ± 6.1%; *Abhd2^−/−^* mothers: 46.7% ± 1.9% of days in estrus), while wild-types had only 35.6% ±5.6% of days in estrus. These results indicate that the observed phenotype was mainly due to the genotype of the pups and not due to epigenetic influence.

### Increased number of atretic antral follicles in *Abhd2^+/−^* and *Abhd2^−/−^*ovaries

To determine whether or not the increased number of ovulated eggs produced during superovulation was due to an altered follicular development, early antral and antral follicles and the corpora lutea of ovaries from *Abhd2^+/+^, Abhd2^+/−^*, and *Abhd2^−/−^* littermate females in proestrus were counted. There was a significant decrease in the number of early antral follicles of both *Abhd2^+/−^* and *Abhd2^−/−^* ovaries compared to those of *Abhd2^+/+^* mice (Fig. 4a). Instead, the females showed an increased number of antral, preovulatory, follicles and, in the case of *Abhd2^+/−^* females, a higher number of corpora lutea (Fig. 4a). However, these differences did not lead to an altered weight of the ovaries when comparing the different genotypes (Supplemental Fig. S3b). Hematoxylin staining of the ovary clearly showed a difference in atretic versus healthy, living follicles, with larger number of atretic follicles present in *Abhd2^+/−^* and *Abhd2^−/−^* ovaries compared to those of *Abhd2^+/+^* mice (Fig. 4b-c). This was further confirmed by terminal deoxynucleotidyl transferase dUTP nick end labeling (TUNEL) staining, where follicles going through atresia showed increased labeling of granulosa cells (Fig. 4d-e). When counting only the healthy antral follicles no genotype specific difference in follicle number was observed. Although the *Abhd2^+/−^* mice displayed increased follicle atresia compared to wild-type mice, the number of healthy antral follicles was comparable to that of *Abhd2^+/+^* ovaries (Fig. 4b).

**Fig. 4.**
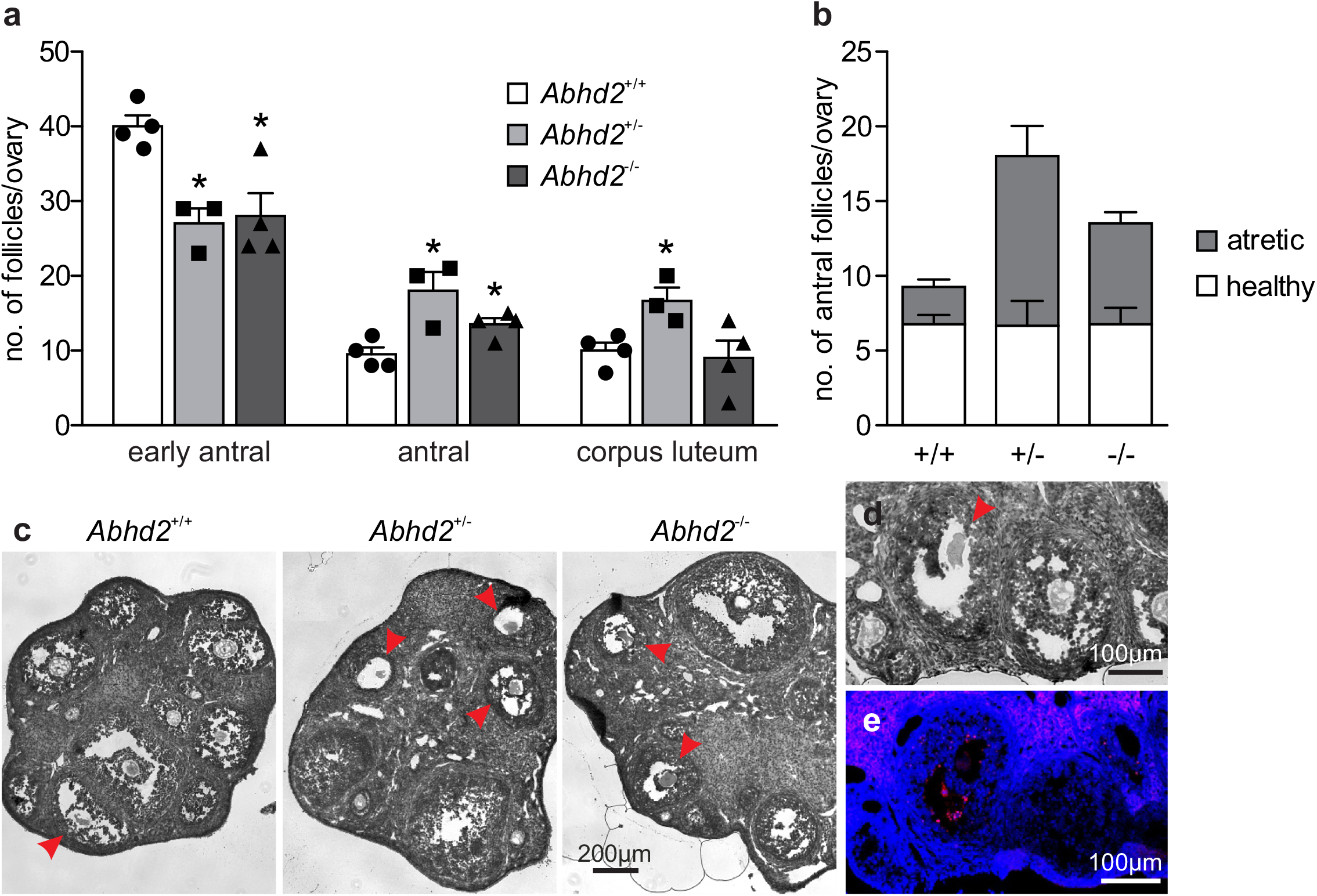
Follicle count of the proestrus ovary. **a)** Ovaries of 3.5-month old *Abhd2^+/−^* and *Abhd2^−/−^* mice showed significantly lower numbers of early antral follicles while the antral follicle number was increased compared to the *Abhd2^+/+^* females. **b)** However, as hematoxylin staining of *Abhd2^+/−^* and *Abhd2^−/−^* ovaries also displayed a higher number of atretic antral follicles compared to the wild-type ovaries, the total number of healthy follicles was similar between genotypes. **c)** Representative images of *Abhd2^+/+^, Abhd2^+/−^* and *Abhd2^−/−^* ovaries with apoptotic antral follicles marked by arrowheads. **d)** and **e)** TUNEL staining confirmed the follicles as apoptotic by labeling the granulosa cells of the atretic follicles. Statistical significance was calculated using the unpaired t-test and the significance of changes is indicated as follows: *, *p ≤ 0.05*.

### ABHD2 is dispensable for *in vitro* follicle culture and ovulation

The increased transition from early antral to antral follicles after *Abhd2* ablation could account for the observed phenotype of superovulated *Abhd2^+/−^* and *Abhd2^−/−^* females. However, as progesterone is needed to initiate the signaling pathways leading to ovulation, *in vitro* follicle culture was performed to determine the role of ABHD2 in this process. Follicles from immature mice were collected and cultured for four days, after which ovulation was induced by addition of hCG to the culture media (Fig. 5a-c). The percentage of ovulated follicles was similar between *Abhd2^+/+^* and *Abhd2^−/−^* females (Fig. 5a). However, as the initial number of large follicles (240 ± 10 μm in diameter) collected from the *Abhd2^−/−^* ovaries was higher than from wild-type ova, the total number of ovulated follicles was increased in cultures from *Abhd2^−/−^* females (Fig. 5b). These results are similar to the higher number of ovulated eggs observed after superovulation of *Abhd2^−/−^* mice. In conclusion, *Abhd2* ablation does not appear to affect the ovulation process directly but rather gives rise to an increased number of mature follicles, possibly by shortening the timing of the diestrus and proestrus stage.

**Fig. 5.**
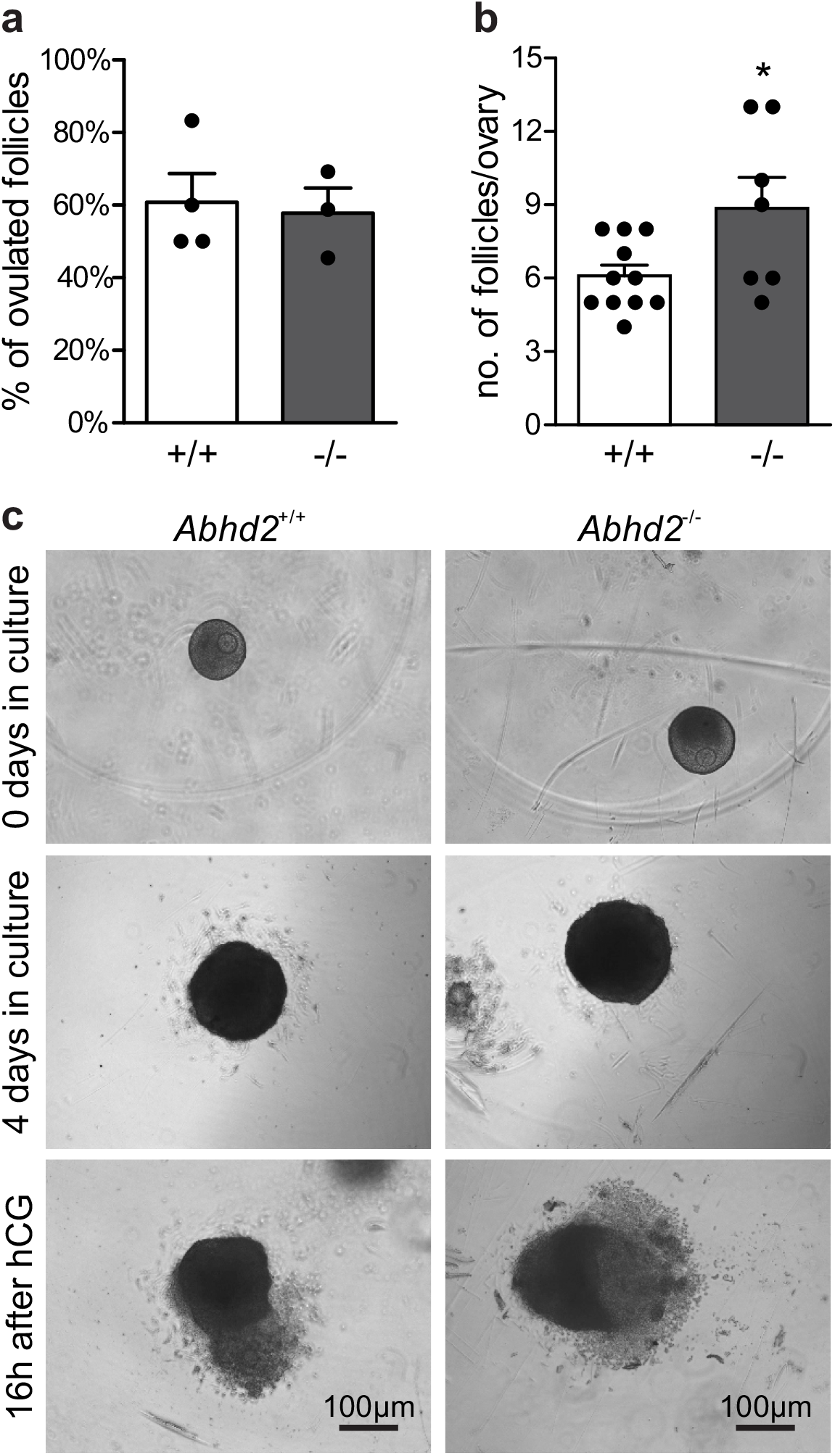
In vitro ovulation. Follicles collected from immature mouse ovaries were cultured to antral stage, after which ovulation was induced by incubation with hCG. **a)** No difference in ovulation efficiency was observed between *Abhd2^+/+^* and *Abhd2^−/−^* mice although **b)** the ovaries of *Abhd2^−/−^* mice contained a higher number of large follicles. **c)** Representative image of *in vitro* ovulation of an *Abhd2^+/+^* and an *Abhd2^−/−^* follicle. Statistical significance was calculated using the unpaired t-test and the significance of changes is indicated as follows: *, *p ≤ 0.05*.

### Changes in ovarian gene expression after *Abhd2* ablation

The observed changes in follicle development and estrous cycle of *Abhd2^−/−^* females might not only result from an altered progesterone signaling but could also be influenced by a change in hormone synthesis. Ovarian hormone production is mainly governed by gonadotropins released by the pituitary gland in the brain. Although mRNA of *Abhd2* has been detected in different brain regions, we have not been able to show the presence of ABHD2 in areas other than the epithelial cells of the choroid plexus (Supplemental Fig. S4), thus making it highly unlikely that the observed ovarian phenotype would be caused by changes in neuronal function. To study the effect of *Abhd2* ablation on steroidogenesis in the ovaries, qPCR was performed to detect expression levels of enzymes involved in hormone synthesis. Ovaries of superovulated *Abhd2^−/−^* females showed no difference in expression of the follicle-stimulating hormone receptor (*Fshr*), a protein vital for follicle development and hormone synthesis, compared to ovaries of *Abhd2^+/+^* mice (Fig. 6a). Neither was there a difference in the expression of cytochrome P450 family members (*Cyp11a1, Cyp17a1* or *Cyp19a1*), the enzymes required for synthesis of progesterone, androgens and estrogens, respectively. Since increased vascularization of the ovary precedes ovulation, the expression of vascular endothelial growth factor A (*Vegfa*), the factor required for vascular endothelial proliferation was also determined. However, no difference in expression between *Abhd2^−/−^* and *Abhd2^+/+^* was detected (Fig. 6a). The expression of nerve growth factor (*Ngf*) and its receptor (nerve growth factor receptor (*Ngfr*)) that are involved in ovulation and in the estrous cycle, were not different between *Abhd2^−/−^* and *Abhd^+/+^* females. However, the expression of the NGF receptor neurotrophic receptor tyrosine kinase 1 (*Ntrk1*, previously known as Tropomyosin receptor kinase A (*TrkA*)) was significantly increased in knockout ovaries compared to those of wild-type mice (Fig. 6b). It is interesting to note that the expression of NGF in stromal cells of the rodent ovary resembles that of ABHD2^15^. Furthermore, increased NGF levels cause the mice to remain in the estrus stage even though they are still cycling^16,17^. NGF signaling is also elevated in women with Polycystic Ovary Syndrome (PCOS)^16,18^ while increased NGF signaling causes mice to show signs of polycystic ovarian morphology (PCOM), with a higher number of atretic antral follicles, without displaying cyst formation- the phenotype we also observe in *Abhd2^−/−^* females.

**Fig. 6.**
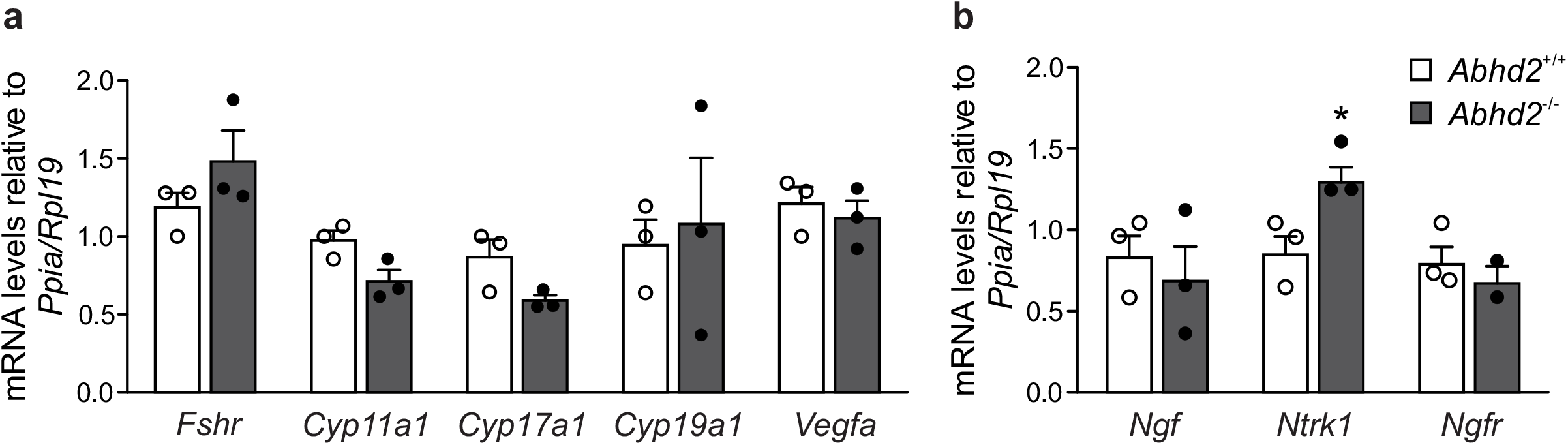
Ovarian gene expression. Relative expression levels of genes involved in steroid synthesis, follicle development and ovulation were measured by qPCR using ovaries of three *Abhd2^+/+^* and three *Abhd2^−/−^* one-month-old superovulated mice. **a)** Expression levels were normalized to the expression of ribosomal protein L19 (*Rpl19*) and peptidylprolyl isomerase A (*Ppia*). Follicle stimulating hormone receptor (*Fshr*), cytochrome P450 family 11 subfamily A member 1 (*Cyp11a1*), cytochrome P450 family 17 subfamily A member 1 (*Cyp17a1*), cytochrome P450 family 19 subfamily A member 1 (*Cyp19a1*), vascular endothelial growth factor A (*Vegfa*), and **b)** nerve growth factor (*Ngf*), neurotrophic receptor tyrosine kinase 1 (*Ntrk1*), and nerve growth factor receptor (*Ngfr*). Statistical significance was calculated using the unpaired t-test and the significance of changes is indicated as follows: *, p ≤ 0.05.

## Discussion

The data presented here shows a novel regulatory role for the membrane progesterone receptor ABHD2 in follicle development and the sexual cycle of females. ABHD2 is already present in pre-pubertal ovaries and the expression remains high in older animals, with localization of the protein to stromal cells surrounding the developing follicles and in corpora lutea. *Abhd2* ablation caused a dysregulation of the estrous cycle rhythm with females remaining in the estrus stage for a longer time while also displaying an increased transition of early antral follicles into pre-ovulatory antral follicles. Injection of PMSG and hCG (which function in a similar manner as the endogenously produced follicle-stimulating hormone (FSH) and luteinizing hormone (LH), respectively^13^) allowed the follicles to develop fully and resulted in the *Abhd2^−/−^* mice ovulating a high number of mature eggs. However, in the native state, the antral follicles displayed signs of atresia and the *Abhd2^−/−^* females ovulated a similar number of eggs as their wild type littermates. ABHD2 was further shown to only regulate follicle maturation and not influence ovulation directly, by *in vitro* culture, where follicles from *Abhd2^−/−^* ovaries responded to hormones in a similar way as those from *Abhd2^+/+^* mice.

Interestingly, the ovarian morphology of *Abhd2^−/−^* mice resemble that of polycystic ovary morphology (PCOM), where the increased number of anovulatory antral follicles can be rescued through administration of FSH^19,20^. This phenotype was described in several mouse models of PCOS where administration of high levels of androgens gives rise to a similar, albeit more severe, dysregulation of ovarian functions as that observed in *Abhd2^−/−^* females. Injection of dehydroepiandrosterone (DHEA) to BALBc females for 20 days caused increased serum levels of both estrogen and progesterone that resulted in the animals remaining in a constant stage of estrus while being unable to ovulate^21,22^. Other mouse models with PCOS also showed anovulation but with the mice estrous cycle remaining in diestrus instead of estrus. Injection of dihydrotestosterone (DHT) to C57Bl/6J females for 90 days led to significantly lower levels of progesterone and a fixed cycle at a so called pseudo-diestrus stage^23^. Balanced steroid secretion and signaling is thus what causes the transition from one stage of the estrous cycle to the next and would implicate a lack in hormone balance of *Abhd2^−/−^* mice. However, the *Abhd2^−/−^* mice did not display a fully developed PCOS as they were able to ovulate, showed similar fertility as wild-type mice in normal mating and did not develop follicular cysts. Furthermore, the fertility rate would indicate that there were no major changes in luteal function or progesterone levels, as the developing embryos were able to implant and mature to full-term. In addition, *Abhd2^−/−^* females still showed estrous cycles with the only differences being a prolonged estrus stage, and shortened diestrus and proestrus stages. Excess hormone exposure during fetal development can also lead to a PCOS phenotype in rodents and results in irregular estrous cycles of the pups^14^. However, we could not detect any difference in estrous cycles when comparing heterozygous mice born from *Abhd2^+/+^, Abhd2^+/−^* or *Abhd2^−/−^* females, further indicating that the hormone levels of the *Abhd2^−/−^* mothers would not be altered in such a way that it could affect the fertility of their offspring. In both rodents and humans, PCOS is often accompanied by a change in insulin resistance and increased body mass index^24,25^. However, *Abhd2^−/−^* females showed regular reduction in glucose levels in glucose tolerance tests and their body weight was not increased compared to *Abhd2^+/+^* females (Supplemental Fig. S3a, c and d). In regards to hormone production there was no significant increase or decrease in the expression of enzymes regulating steroid production in *Abhd2^−/−^* ovaries after superovulation, which further supports a phenotype of mildly altered or normal steroid levels in *Abhd2^−/−^* mice.

The only detected difference in gene expression after superovulation of wild-type and *Abhd2^−/−^* mice was an increased expression of the NGF receptor *Ntrk1*. As previously mentioned, the expression of NGF in stromal cells of the rodent ovary resembles that of ABHD2^15^. However, similar to ABHD2, NGF signaling does not only regulate the function of the interstitial cells but can also affect the maturation of granulosa cells, as seen in a mouse model with excess expression of *Ngf*. The *Ngf* overexpressors present with increased antral formation and apoptosis of granulosa cells, which leads to a PCOM phenotype without the formation of ovarian cysts^16,17^. However, an additional, mild increase in LH-levels caused cysts to form, which would indicate that increased NGF signaling could promote the formation of PCOS^16^. Thus, further studies are needed to determine if a change in pituitary hormone secretion could lead to a more prominent phenotype of *Abhd2* KO mice.

The estrous cycle of mice also changes during aging, with fertile animals spending more time in estrus than older animals who usually become acyclic and remain in diestrus^26^. This was shown in a mouse model of premature ovarian failure where the tumor necrosis receptor type I (*Tnfr1*) was ablated. Younger *Tnfr1* knockout females, similar to the *Abhd2^−/−^* mice, displayed a stronger ovulatory response to hormone injections than wild-type females. Young animals also showed a longer period in estrus which was similar to estrous cycles of 6-month-old females^27^. The phenotype was caused by a premature sexual maturity which led to an early onset of senescence at 6 months of age with lower number of litters produced and a disruption in estrous cyclicity^27^. Similarly, a mouse model with granulosa cell specific ablation of neuregulin 1 (*Nrg1*) showed signs of early ovarian failure where the estrous cycle stayed in a continuous, so called, weak estrus stage^28^. These mice also displayed increased fibrosis of ovarian stroma and lower numbers of progeny, a phenotype similar to that of older animals^28^. The premature ovarian aging of both *Nrg1* and *Tnfr1* knockout females is related to a reduced response to FSH and LH^27,28^. Although *Abhd2* KO females show a prolonged estrus stage at around 3 months of age, this did not lead to a premature decline in fertility at 8 months of age, as was determined by breeding data from *Abhd2^+/−^* and *Abhd2^−/−^* females. In addition, there was no observed early depletion in ovulated follicles due to the increased number of antral follicles of *Abhd2* KO females. This was shown after superovulation of older *Abhd2^+/−^* and *Abhd2^−/−^* females which still produced similar number of eggs compared to wild-type mice of similar age. It is more likely that the transition from early antral to antral follicle differentiation was altered, as the higher number of antral follicles correlated with a similar decrease in number of early antral follicles. Such a transition might not be enough to give rise to depletion of the ovarian follicle reserve. Instead we propose that the increased number of anovulatory antral follicles at proestrus could cause the prolonged estrus stage observed in *Abhd2^−/−^* females. The mice would remain in estrus until the follicles have gone through apoptosis and would thereafter proceed toward the next stage of the estrous cycle. The lack of membrane progesterone receptor (ABHD2) in stromal cells at diestrus, the stage with highest progesterone production^29^, could then cause the cycle to move rapidly into the follicular phase.

In humans, downregulation of *ABHD2* expression resulted in anoikis resistance in high-grade serous ovarian cancer^30^. The phenotype was attributed to an increased phosphorylation of p38MAPK and MAPK3/1, the signaling pathways which have previously been shown to regulate anoikis resistance^30–32^. In healthy ovaries MAPK3/1 (also known as ERK1/2) signaling is needed for multiple steps in follicle maturation. In early antral follicles MAPK3/1 phosphorylation inhibits proliferation and induces differentiation of granulosa cells, while in antral follicles it stimulates cumulus cell expansion and ovulation^33^. Considering the phenotype of *Abhd2^−/−^* mice, an increased MAPK3/1 signaling could cause the observed increase in antral follicle differentiation while further activation through LH signaling could give rise to ovulation of the mature follicles.

In conclusion, *Abhd2* ablation caused an altered follicle maturation which led to a higher number of antral follicles with an atretic phenotype. In addition, a change in estrous cyclicity was observed with mice spending longer times in estrus. Future studies will focus on the functional role of ABHD2 in this process, which could affect several progesterone signaling pathways in the ovary, including phosphorylation of MAPK/ERK. Furthermore, because of a phenotype resembling PCOS and the known expression of ABHD2 in human ovaries, continued studies of the role of ABHD2 in human follicle maturation and the menstrual cycle could be vital to explain the disease process of PCOS.

## Methods

### Ethics statement

All mice were kept in the Animal Facility of the University of California, Berkeley, fed standard chow diet (PicoLab Rodent diet 20, #5053, LabDiet) and hyper-chlorinated water *ad libitum* in a room with controlled light (14 hours light, 10 hours darkness) and temperature (23±0.5°C). Animals were humanely killed according to ACUC guidelines with every effort made to minimize suffering. All experiments were performed in accordance with NIH Guidelines for Animal Research and approved by UC Berkeley Animal Care and Use Committee (AUP 2015-07-7742).

### Generation of *Abhd2* KO mice using CRISPR Ribonucleoprotein Electroporation of Zygotes (CRISPR-EZ)

The CRISPR-EZ method, where single guide RNA (sgRNA)/Cas9 complexes are delivered into mouse zygotes by electroporation, was developed by Chen et al and Modzelewski et al 11,12.

### In Vitro Synthesis of sgRNAs

SgRNAs to target the introns flanking *Abhd2* exon 6 were designed as previously described. Briefly, we utilized the Gene Perturbation Platform^34^, Chop-Chop^35^, and CRISPR Design^36^ algorithms to design sgRNAs for the target sequence. The obtained 20-nt sequences were incorporated into a DNA oligonucleotide template containing a T7 promoter and a sgRNA scaffold by overlapping PCR using Phusion high fidelity DNA polymerase (New England Biolabs, M0530). The primer sequences for overlapping PCR were as previously described^11^ including the designed sgRNA sequences; sgRNA1: 5’-TGTGCTGAGAAAACGCAGGT-3’ and sgRNA2: 5’-TGTCAACTGAAGGACCCAAG-3’) The template sequence was transcribed into RNA using a T7 RNA polymerase (New England Biolabs, E2040S) after which the DNA template was removed by treatment with RNase-Free DNaseI (New England Biolabs, M0303S). The produced sgRNAs were further purified by allowing them to bind to SeraMeg Speedbeads magnetic carboxylate-modified particles (GE Healthcare, 65152105050250. After washing the bead/RNA pellets with 80% ethanol the sgRNAs were eluted in nuclease free water (Ambion, AM9937) and stored at −80 °C until use.

### Delivery of sgRNA/Cas9 complexes into zygotes

To form a ribonucleoprotein complex, the Cas9 protein (QB3 Macrolab, University of California at Berkeley) was incubated with the two sgRNAs in a solution containing 20 mM HEPES (Sigma, H7523), pH 7.5, 150 mM KCl (Fisher Chemical, P217), 1 mM MgCl_2_ (Sigma, 68475), 10% glycerol, (Sigma, G2025) and 1 mM reducing agent DL-Dithiothreitol (Sigma, D0632), to a final molar ratio of 1:2. The ribonucleoprotein mixture was prepared at 37 °C for 10 min immediately before electroporation. To obtain zygotes, four-week-old C57Bl/6N (Charles River) female mice were superovulated by intraperitoneal injection (i.p.) of 5 IU pregnant mare serum gonadotropin (PMSG, Sigma, G4877) and 48 hours later, 5 IU human chorionic gonadotropin (hCG, Millipore, 230734), after which they were immediately put in mating with fertile 2-4-month old C57Bl/6N males. 12 hours post-coitum, females with a plug were euthanized and the one-cell zygotes were collected from the ampulla of the oviduct. The cumulus cells surrounding the fertilized eggs were removed by hyaluronidase treatment (Life Global Group, LGHY-010) and the zona pellucida was weakened by incubation in acidic Tyrode’s solution (Sigma, T1788) as previously described^11^. For delivery of the sgRNA/Cas9 complex, ~40 zygotes were pooled in Opti-MEM reduced serum media (Gibco, 31985-070) containing the pre-formed Cas9/sgRNA ribonucleoproteins and loaded into a 1-mm electroporation cuvette (BioRad, 165-2089). Electroporation was performed using the GenePulser Xcell (BioRad, 1652660) with six pulses at 30 V for 3 ms, separated by a 100 ms interval. Immediately after electroporation the zygotes were washed in KSOM + AA media (KCl-enriched simplex optimization medium with amino acid supplement, Zenith Biotech, ZEKS-050) supplemented with 4 mg/ml bovine serum albumin (BSA, Sigma, A9647) and cultured overnight in KSOM/BSA at 37°C, 5% CO_2_. The following day, the embryos that had developed into two-cell stage were implanted into pseudopregnant CD-1 females (Envigo) at ~10 embryos per oviduct by the UC Berkeley Cancer Research Laboratory, Gene Targeting Facility. The resulting progeny was genotyped using EmeraldAmp GT PCR Master Mix (Takara, RR310) and primers flanking the deleted region of *Abhd2*, Abhd2 Fw: 5’- AGGGCTTAACTCTTGCTGGT-3’, Abhd2 Rev: 5’-ACTCAGACACGATCCGAGAC-3’, Tm: 56°C. Each mouse, positive for the deleted region of *Abhd2*, was placed in breeding with a C57Bl/6N mouse of the opposite gender. The heterozygous progeny of these matings were further bred to produce *Abhd2^−/−^* mice and the control *Abhd2^+/+^* littermates.

### Breeding efficiency and spontaneous ovulation

To determine the breeding efficiency of *Abhd2^+/+^, Abhd2^+/−^*, and *Abhd2^−/−^* female mice, two-month-old females of each genotype were placed in breeding with either *Abhd2^+/+^* males (*Abhd2^+/+^* and *Abhd2^−/−^* females) or *Abhd2^+/−^* males (*Abhd2^+/−^* and *Abhd2^−/−^* females) for ~6 months. Two or more breeding pairs of each genotype combination were used to determine the average number of offspring per litter and the birth rate of female mice. Further, to detect the number of spontaneously ovulated eggs, two-three-month-old females in proestrus were placed in breeding with fertile *Abhd2^+/+^* male mice. When a copulatory plug was detected, the females were euthanized and the fertilized zygotes were collected from the oviduct ampulla in Dulbecco’s phosphate buffered saline (DPBS, Gibco, 14190-144) and counted.

### Superovulation and *in vitro* fertilization

Sperm from the cauda epididymides of 2-month-old C57Bl/6N male mice were allowed to swim out in Embryomax Human Tubal Fluid medium (HTF, Millipore, MR-070-D) and incubated for capacitation, 1 hour at 37°C, 5% CO_2_. To collect eggs from seven *Abhd2^+/+^*, eight *Abhd2^+/−^*, and seven *Abhd2^−/−^* females, superovulation of 3.5-4 week old mice was performed using i.p. injection of PMSG and hCG as described above. 13 hours after hCG injection the females were euthanized and the cumulus-oocyte complexes were collected from the oviduct ampulla in HTF medium and counted. *In vitro* fertilization (IVF) was performed as previously described^37^ using 300 000 sperm/mL HTF medium. After fertilization the zygotes were washed in KSOM + AA media, supplemented with 4 μl BSA and divided into ~15 zygotes/10 μl KSOM/BSA for culture at 37°C, 5% CO_2_. 3.5 days following IVF, the number of fertilized eggs was determined as the percentage of embryos that had reached morula or blastula stage.

### Evaluation of the mouse estrous cycle

Each stage of the mouse estrous cycle was determined in the afternoon by cytological assessment of vaginal smear samples. Samples were collected from 2.5-month-old *Abhd2^+/+^*, *Abhd2^+/−^*, and *Abhd2^−/−^* mice, three females of each genotype, every day for one month. DPBS was used to collect samples of each mouse, using the methods previously described^38^. Samples were separately placed on glass slides and examined under an Olympus IX-75 inverted light microscope for cellular contents. Visual representation of each estrous cycle phase indicated by Zenclussen et al. was used as a reference^39^.

### Ovarian follicle count

To count the total number of follicles in *Abhd2^+/+^, Abhd2^+/−^*, and *Abhd2^−/−^* ovaries, 3-4 3.5-month-old female mice of each genotype were euthanized at proestrus stage, and the ovaries were dissected out and weighed. The larger of the two ovaries was fixed in 4% paraformaldehyde (Electron Microscopy Sciences, 15714), embedded in optimal cutting temperature compound (OCT, Sakura, 4583), frozen on dry ice, and stored at −80°C until use. The ovaries were sectioned through and every fifth section of 8μm was mounted on glass slides and stained with hematoxylin 1 (Fisher HealthCare, 220-101) using standard procedures. Images of each ovarian section were taken at 50× magnification and organized sequentially for follicle counting. Using this method, we were able to only count each follicle once, even though the larger follicles were present in several sections. Each follicle was categorized as early antral, antral, or corpus luteum based on their morphology. Early antral follicles were identified by the presence of segmented cavities between multiple layers of granulosa cells surrounding the oocyte. Antral follicles were identified by the presence of a large continuous antral cavity, and corpus luteum by the absence of oocytes. Atretic antral follicles were defined by a thinning of the granulosa cell layer, which displayed many pyknotic cells, and the additional lack of a clear cumulus cell layer surrounding the oocyte. The morphology of the atretic antral follicles were further confirmed by comparing hematoxylin and TUNEL stainings.

### *In vitro* ovulation

Ovaries were retrieved from eleven 20-to 21-day-old wild-type (C57Bl/6N) or seven *Abhd2^−/−^* female mice as described previously^40,41^. Briefly, ovaries were placed in dissection media (L15 media (Gibco, 11415-064), 0.5% PenStrep (Fisher Scientific, 15140-122) and 1% fetal bovine serum (FBS, X&Y Cell Culture, FBS-500-HI)), and the bursal sac and any excess material were removed. Each ovary was halved and incubated in collagenase media (Minimum essential medium alpha (αMEM, Gibco, 12561-056), 0.5% PenStrep, 0.8% type I collagenase (Worthington, LS004194), and 1% DNase I (Fisher Scientific, EN0521) at 37°C and 5% CO_2_ for 40 min. Preantral follicles were mechanically isolated using 28 gauge insulin needles in dissection media. Only intact follicles that were 240 ± 10 μm in diameter were isolated and incubated at 37°C and 5% CO_2_ in maintenance media (αMEM, 0.5% PenStrep, and 1% FBS) for 2 hours before encapsulation. Follicles were encapsulated in 0.7% alginate and cultured as previously described^40,41^, with slight modifications. The encapsulated follicles were placed individually in a 96-well plate (Fisher Scientific, 08-772-53) and cultured for four days in 100 μL growth media (αMEM, 3 mg/mL BSA, 1 mg/mL bovine fetuin (Sigma, F3385), 0.01 IU/mL recombinant human FSH (NHPP, Dr. Parlow), 5 μg/mL insulin, 5 μg/mL transferrin, and 5 ng/mL selenium (Sigma, I1884)). At day 2 of culture, half of the growth media was exchanged. After 4 days in culture, the alginate was removed by incubating the follicles in alginate lyase media (αMEM, 0.5% PenStrep, 1% FBS, and 1mg/mL alginate lyase (Sigma, A1603)) at 37°C and 5% CO_2_ for 30 minutes. Thereafter, antral follicles larger than 330 ± 15 μm in diameter were washed in ovulation media (αMEM, 10% FBS, 5 ng/mL EGF (Sigma, E4127), 1.5 IU/mL hCG, 5 μg/mL insulin, 5 μg/mL transferrin, and 5 ng/mL selenium) and placed individually in a 96-well plate containing 100 μL of ovulation media per well. After 16 hours at 37°C and 5% CO_2_ the percentage of successful ovulation was calculated as the number of follicles with a clear visual evidence of follicle rupture and oocyte extrusion compared to total follicle number.

### Western blotting

To detect ABHD2 protein in ovaries and different brain regions, 3.5-4-week-old *Abhd2^+/+^, Abhd2^+/−^*, and *Abhd2^−/−^* superovulated mice and a 2.5-month-old *Abhd2^+/+^* female respectively, were euthanized, the tissue dissected and snap frozen, and protein was isolated using standard procedures. The samples were analyzed by Western blotting, using a rabbit polyclonal anti-ABHD2 antibody (1:10 000 dilution, Proteintech Group, 14039-1-AP) and a peroxidase-conjugated anti-rabbit secondary antibody (1:15000 dilution, Abcam, ab6721). To ensure equal sample loading, the membrane was stripped by incubation with OneMinute Plus Strip (GM Biosciences, GM6011) according to the manufacturer’s instructions. Thereafter, the membrane was re-hybridized with a mouse monoclonal antiactin antibody (1:5000 dilution, Abcam, ab3280-500) and a peroxidase-conjugated goat anti-mouse secondary antibody (1:15000 dilution, EMD-Millipore, AP181P).

### Immunohistochemistry

Ovaries of superovulated 3.5-4 week-old *Abhd2^+/+^* and *Abhd2^−/−^* mice were fixed overnight at 4°C in 4% PFA, and the ovary of a 1-week-old *Abhd2^+/+^* female was fixed for 4 hours after which they went through a sucrose gradient (10%-20%-30% sucrose/PBS) and were frozen in OCT. To detect localization of ABHD2 in the ovary, 8 μm sections were placed on charged slides and immediately used for staining, according to standard procedures. In brief, for antigen retrieval the sections were incubated in 1% SDS solution for 5 minutes, blocked in 5% BSA for 1 hour at room temperature and incubated overnight at 4°C with rabbit polyclonal anti-ABHD2 (1:300) and mouse monoclonal anti-α-actin (1:500, sc-32251, Santa Cruz Biotechnology). The antibody-antigen complexes were visualized by incubation for 1 hour at room temperature with 1:2000 Alexa Fluor 488-conjugated goat anti-rabbit (Molecular Probes A11008/Jackson ImmunoResearch, 111-485-144) and Cy5-conjugated donkey anti-mouse (Jackson ImmunoResearch, 715-175-150) or anti-rabbit (Jackson ImmunoResearch, 711-015-152) antibodies. The sections were mounted using ProLong Gold antifade reagent with DAPI (Invitrogen, P36935) and imaged using confocal laser scanning microscopy (Olympus Fluoview FV1000).

### Reverse transcriptase and quantitative RT-PCR

For analysis of gene expression, 21- and 28-day old *Abhd2^+/+^* mice were euthanized and the ovaries dissected out and snap frozen. Samples from superovulated 1-month-old mice, and 2-month-old mice at proestrus and estrus, were similarly collected and frozen. Total RNA was isolated from the tissues using TRIzol Reagent according to the manufacturer’s instructions (Ambion, 15596018). For reverse transcription, 1 μg of total RNA was treated with DNaseI and reverse-transcribed by using the RevertAid H Minus RT enzyme (Fisher Scientific, EP0451). The cDNA was diluted 1:50–1:100 for qPCR. qPCR was performed using the DyNAmo HS SYBR Green qPCR Master Mix (Fisher Scientific, F-410). All samples were run in triplicate reactions. For analysis of *Abhd2* expression at different time points in wild-type mice ovaries primers binding to exon 3 and exon 4 were used (*Abhd2* e3 Fw and *Abhd2* e4 Re). To determine expression of the deleted exon 6 in *Abhd2* KO mice primers binding in exon 5 and exon 6 were used (*Abhd2* e5 Fw and *Abhd2* e6 Re). Ribosomal protein L19 (*Rpl19*) and peptidylprolyl isomerase A (*Ppia*) were used as endogenous controls to equalize for the amounts of RNA in the ovaries. Primer sequences for *Rpl19, Ppia*, and cytochrome P450 family 19 subfamily A member 1 (*Cyp19a1*) were as previously described^42,43^. Primer sequences and qPCR conditions for analyzing the expression of *Abhd2, Fshr*, cytochrome P450 family 11 subfamily A member 1 (*Cyp11a1*), cytochrome P450 family 17 subfamily A member 1 (*Cyp17a1*), *Vegfa, Ngf, Ntrk1*, and *Ngfr* are described in Supplemental Table 1.

### Intraperitoneal glucose tolerance test

To detect the effect of *Abhd2* ablation on glucose uptake, 2.5-month-old *Abhd2^+/+^* (n=5), *Abhd2^+/−^* (n=7), and *Abhd2^−/−^* (n=6) female mice were fasted overnight (16 hours). Blood glucose levels were measured as mmol/l by tail vein sampling using Contour Next ONE meter (Ascensia Diabetes Care) and Contour Next blood glucose test strips (Ascensia Diabetes Care). Immediately after baseline measurement (0 min) the mice were given an i.p. injection of 2 g/kg glucose (20% solution) and blood glucose levels were measured at 15, 30, 60 and 120 min after injection.

### Statistical analyses

For statistical analyses of fertility efficiency, ovulation number, days in estrus, follicle numbers, and gene expression levels the GraphPad Prism 5 software (GraphPad Software, Inc., La Jolla, CA) was used. Unpaired t-test was used to determine statistical significance, assigning *p ≤ 0.05* as the limit. All results are shown with standard error of mean.

## Acknowledgments

This work was supported by R01GM111802, Pew Biomedical Scholars Award and Packer Wentz Endowment Will (to PVL). We thank S. Chen (Department of Molecular and Cell Biology, University of California, Berkeley) for preparation of the SeraMeg SpeedBeads for sgRNA purification and A. Shikanov (Department of Biomedical Engineering, University of Michigan) for providing the protocol for *in vitro* follicle culture.

## Author contributions

IB and PVL conceived the project, designed the experiments and wrote the manuscript. IB performed, or assisted in the completion, of all studies, data acquisition and analysis for the manuscript. DHC collected samples for estrous stage analysis, determined the follicle count and performed *in vitro* follicle culture. SM assisted in maintenance of the mouse colony and performing molecular biology analysis. LG assisted in performing the *in vitro* fertilization studies. AM provided assistance in designing the sgRNAs and assisted with performing the zygote electroporation. LH helped with CRISPR-EZ by providing essential reagents and support. All authors discussed the results and commented on the manuscript.

## Declaration of Interests

LH is an inventor on patents of an electroporation-based CRISPR technology for mouse genome engineering and is a co-founder of a company to further develop this technology for mammalian genome editing.

## Notes

#### Summary of Updates

Author name updated

## References

1 Roy, S. K. & Greenwald, G. S. In vitro steroidogenesis by primary to antral follicles in the hamster during the periovulatory period: effects of follicle-stimulating hormone, luteinizing hormone, and prolactin. Biol Reprod 37, 39–46, doi:10.1095/biolreprod37.1.39 (1987).

2 Hashimoto-Partyka, M. K., Lydon, J. P. & Iruela-Arispe, M. L. Generation of a mouse for conditional excision of progesterone receptor. Genesis 44, 391–395, doi:10.1002/dvg.20227 (2006).

3 Lydon, J. P. et al. Mice lacking progesterone receptor exhibit pleiotropic reproductive abnormalities. Genes & development 9, 2266–2278, doi:10.1101/gad.9.18.2266 (1995).

4 Walmer, D. K., Wrona, M. A., Hughes, C. L. & Nelson, K. G. Lactoferrin expression in the mouse reproductive tract during the natural estrous cycle: correlation with circulating estradiol and progesterone. Endocrinology 131, 1458–1466, doi:10.1210/endo.131.3.1505477 (1992).

5 Revelli, A., Massobrio, M. & Tesarik, J. Nongenomic actions of steroid hormones in reproductive tissues. Endocr Rev 19, 3–17 (1998).

6 Baldi, E. et al. Intracellular calcium accumulation and responsiveness to progesterone in capacitating human spermatozoa. J Androl 12, 323–330 (1991).

7 Blackmore, P. F., Beebe, S. J., Danforth, D. R. & Alexander, N. Progesterone and 17 alpha-hydroxyprogesterone. Novel stimulators of calcium influx in human sperm. J Biol Chem 265, 1376–1380 (1990).

8 Blackmore, P. F., Neulen, J., Lattanzio, F. & Beebe, S. J. Cell surface-binding sites for progesterone mediate calcium uptake in human sperm. J Biol Chem 266, 18655–18659 (1991).

9 Poisbeau, P. et al. Analgesic strategies aimed at stimulating the endogenous production of allopregnanolone. Frontiers in cellular neuroscience 8, 174, doi:10.3389/fncel.2014.00174 (2014).

10 Miller, M. R. et al. Unconventional endocannabinoid signaling governs sperm activation via the sex hormone progesterone. Science 352, 555–559, doi:10.1126/science.aad6887 (2016).

11 Chen, S., Lee, B., Lee, A. Y., Modzelewski, A. J. & He, L. Highly Efficient Mouse Genome Editing by CRISPR Ribonucleoprotein Electroporation of Zygotes. J Biol Chem 291, 14457–14467, doi:10.1074/jbc.M116.733154 (2016).

12 Modzelewski, A. J. et al. Efficient mouse genome engineering by CRISPR-EZ technology. Nature protocols 13, 1253–1274, doi:10.1038/nprot.2018.012 (2018).

13 Christenson, C. M. & Eleftheriou, B. E. Dose-dependence of superovulation response in mice to two injections of PMSG. J Reprod Fertil 29, 287–289 (1972).

14 Abbott, D. H., Padmanabhan, V. & Dumesic, D. A. Contributions of androgen and estrogen to fetal programming of ovarian dysfunction. Reprod Biol Endocrinol 4, 17, doi:10.1186/1477-7827-4-17 (2006).

15 Dissen, G. A. et al. A role for trkA nerve growth factor receptors in mammalian ovulation. Endocrinology 137, 198–209, doi:10.1210/endo.137.1.8536613 (1996).

16 Dissen, G. A. et al. Excessive ovarian production of nerve growth factor facilitates development of cystic ovarian morphology in mice and is a feature of polycystic ovarian syndrome in humans. Endocrinology 150, 2906–2914, doi:10.1210/en.2008-1575 (2009).

17 Wilson, J. L. et al. Excess of nerve growth factor in the ovary causes a polycystic ovary-like syndrome in mice, which closely resembles both reproductive and metabolic aspects of the human syndrome. Endocrinology 155, 4494–4506, doi:10.1210/en.2014-1368 (2014).

18 Gulino, F. A., Giuffrida, E., Leonardi, E., Marilli, I. & Palumbo, M. A. Intrafollicular nerve growth factor concentration in patients with polycystic ovary syndrome: a case-control study. Minerva ginecologica 68, 110–116 (2016).

19 Perales-Puchalt, A. & Legro, R. S. Ovulation induction in women with polycystic ovary syndrome. Steroids 78, 767–772, doi:10.1016/j.steroids.2013.05.005 (2013).

20 Tessaro, I. et al. Effect of oral administration of low-dose follicle stimulating hormone on hyperandrogenized mice as a model of polycystic ovary syndrome. Journal of ovarian research 8, 64, doi:10.1186/s13048-015-0192-9 (2015).

21 Luchetti, C. G. et al. Effects of dehydroepiandrosterone on ovarian cystogenesis and immune function. J Reprod Immunol 64, 59–74, doi:10.1016/j.jri.2004.04.002 (2004).

22 Sander, V. et al. Role of the N, N’-dimethylbiguanide metformin in the treatment of female prepuberal BALB/c mice hyperandrogenized with dehydroepiandrosterone. Reproduction 131, 591–602, doi:10.1530/rep.1.00941 (2006).

23 Caldwell, A. S. et al. Characterization of reproductive, metabolic, and endocrine features of polycystic ovary syndrome in female hyperandrogenic mouse models. Endocrinology 155, 3146–3159, doi:10.1210/en.2014-1196 (2014).

24 Sirmans, S. M. & Pate, K. A. Epidemiology, diagnosis, and management of polycystic ovary syndrome. Clinical epidemiology 6, 1–13, doi:10.2147/CLEP.S37559 (2013).

25 van Houten, E. L. & Visser, J. A. Mouse models to study polycystic ovary syndrome: a possible link between metabolism and ovarian function? Reproductive biology 14, 32–43, doi:10.1016/j.repbio.2013.09.007 (2014).

26 Nelson, J. F., Felicio, L. S., Randall, P. K., Sims, C. & Finch, C. E. A longitudinal study of estrous cyclicity in aging C57BL/6J mice: I. Cycle frequency, length and vaginal cytology. Biol Reprod 27, 327–339, doi:10.1095/biolreprod27.2.327 (1982).

27 Roby, K. F., Son, D. S. & Terranova, P. F. Alterations of events related to ovarian function in tumor necrosis factor receptor type I knockout mice. Biol Reprod 61, 1616–1621, doi:10.1095/biolreprod61.6.1616 (1999).

28 Umehara, T. et al. The acceleration of reproductive aging in Nrg1(flox/flox);Cyp19-Cre female mice. Aging cell 16, 1288–1299, doi:10.1111/acel.12662 (2017).

29 MacDonald, J. K., Pyle, W. G., Reitz, C. J. & Howlett, S. E. Cardiac contraction, calcium transients, and myofilament calcium sensitivity fluctuate with the estrous cycle in young adult female mice. American journal of physiology. Heart and circulatory physiology 306, H938–953, doi:10.1152/ajpheart.00730.2013 (2014).

30 Yamanoi, K. et al. Suppression of ABHD2, identified through a functional genomics screen, causes anoikis resistance, chemoresistance and poor prognosis in ovarian cancer. Oncotarget 7, 47620–47636, doi:10.18632/oncotarget.9951 (2016).

31 Cai, Q., Yan, L. & Xu, Y. Anoikis resistance is a critical feature of highly aggressive ovarian cancer cells. Oncogene 34, 3315–3324, doi:10.1038/onc.2014.264 (2015).

32 Carduner, L. et al. Cell cycle arrest or survival signaling through alphav integrins, activation of PKC and ERK1/2 lead to anoikis resistance of ovarian cancer spheroids. Exp Cell Res 320, 329–342, doi:10.1016/j.yexcr.2013.11.011 (2014).

33 Fan, H. Y. et al. MAPK3/1 (ERK1/2) in ovarian granulosa cells are essential for female fertility. Science 324, 938–941, doi:10.1126/science.1171396 (2009).

34 Doench, J. G. et al. Optimized sgRNA design to maximize activity and minimize off-target effects of CRISPR-Cas9. Nature biotechnology 34, 184–191, doi:10.1038/nbt.3437 (2016).

35 Montague, T. G., Cruz, J. M., Gagnon, J. A., Church, G. M. & Valen, E. CHOPCHOP: a CRISPR/Cas9 and TALEN web tool for genome editing. Nucleic acids research 42, W401–407, doi:10.1093/nar/gku410 (2014).

36 Hsu, P. D. et al. DNA targeting specificity of RNA-guided Cas9 nucleases. Nature biotechnology 31, 827-832, doi:10.1038/nbt.2647 (2013).

37 Navarrete, F. A. et al. Biphasic role of calcium in mouse sperm capacitation signaling pathways. J Cell Physiol 230, 1758–1769, doi:10.1002/jcp.24873 (2015).

38 McLean, A. C., Valenzuela, N., Fai, S. & Bennett, S. A. Performing vaginal lavage, crystal violet staining, and vaginal cytological evaluation for mouse estrous cycle staging identification. Journal of visualized experiments: JoVE, e4389, doi:10.3791/4389 (2012).

39 Zenclussen, M. L., Casalis, P. A., Jensen, F., Woidacki, K. & Zenclussen, A. C. Hormonal Fluctuations during the Estrous Cycle Modulate Heme Oxygenase-1 Expression in the Uterus. Frontiers in endocrinology 5, 32, doi:10.3389/fendo.2014.00032 (2014).

40 Skory, R. M., Xu, Y., Shea, L. D. & Woodruff, T. K. Engineering the ovarian cycle using in vitro follicle culture. Hum Reprod 30, 1386–1395, doi:10.1093/humrep/dev052 (2015).

41 Xu, M., Kreeger, P. K., Shea, L. D. & Woodruff, T. K. Tissue-engineered follicles produce live, fertile offspring. Tissue engineering 12, 2739–2746, doi:10.1089/ten.2006.12.2739 (2006).

42 Bjorkgren, I. et al. Dicer1 ablation in the mouse epididymis causes dedifferentiation of the epithelium and imbalance in sex steroid signaling. PLoS One 7, e38457, doi:10.1371/journal.pone.0038457 (2012).

43 Hakkarainen, J. et al. Hydroxysteroid (17beta)-dehydrogenase 1-deficient female mice present with normal puberty onset but are severely subfertile due to a defect in luteinization and progesterone production. FASEB journal: official publication of the Federation of American Societies for Experimental Biology 29, 3806–3816, doi:10.1096/fj.14-269035 (2015).

