## Supplementary Table and Figures for "Alpha/Beta Hydrolase Domain-Containing Protein 2 regulates the rhythm of follicular maturation and estrous stages of the female reproductive cycle"

**Supplemental Table 1**

| Gene | | 5'- primer – 3' sequence | Tm ˚C |
|---|---|---|---|
| abhydrolase domain containing 2 | Abhd2 e3 Fw | CCGTAGCTGCCGTTCTCTAC | 61 |
| | Abhd2 e4 Re | CTCCCCATCTTCCCATACAA | |
| | Abhd2 e5 Fw | GAATTGCCAACCACAGCGAG | 61 |
| | Abhd2 e6 Rev | AGCTGGGTCTGGGGATATGT | |
| follicle stimulating hormone receptor | Fshr Fw | TGTTTTCCAGGGAGCCTCTG | 61 |
| | Fshr Rev | AGCAGTGACTGGGGTAGGTA | |
| cytochrome P450 family 11 subfamily A member 1 | Cyp11a1 Fw | ATTACCGAGATGCTGGCAGG | 60 |
| | Cyp11a1 Rev | GTGTCTCCTTGATGCTGGCT | |
| cytochrome P450 family 17 subfamily A member 1 | Cyp17a1 Fw | CAATGACCGGACTCACCTCC | 60 |
| | Cyp17a1 Rev | CCTTCGGGATGGCAAACTCT | |
| vascular endothelial growth factor A | Vegfa Fw | CGATTGAGACCCTGGTGGAC | 61 |
| | Vegfa Rev | GCTGGCTTTGGTGAGGTTTG | |
| nerve growth factor | Ngf Fw | TGTGCCTCAAGCCAGTGAAA | 60 |
| | Ngf Rev | CACTGAGGTGAGCTTGGGTC | |
| neurotrophic receptor tyrosine kinase 1 | Ntrk1 Fw | CATCGTGCGCTTCTTTGGAG | 60 |
| | Ntrk1 Rev | CAGCAGCTTTGCATCAGGTC | |
| nerve growth factor receptor (TNFR superfamily, member 16) | Ngfr Fw | CCGCTGACAACCTCATTCCT | 60 |
| | Ngfr Rev | TGTCGCTGTGCAGTTTCTCT | |

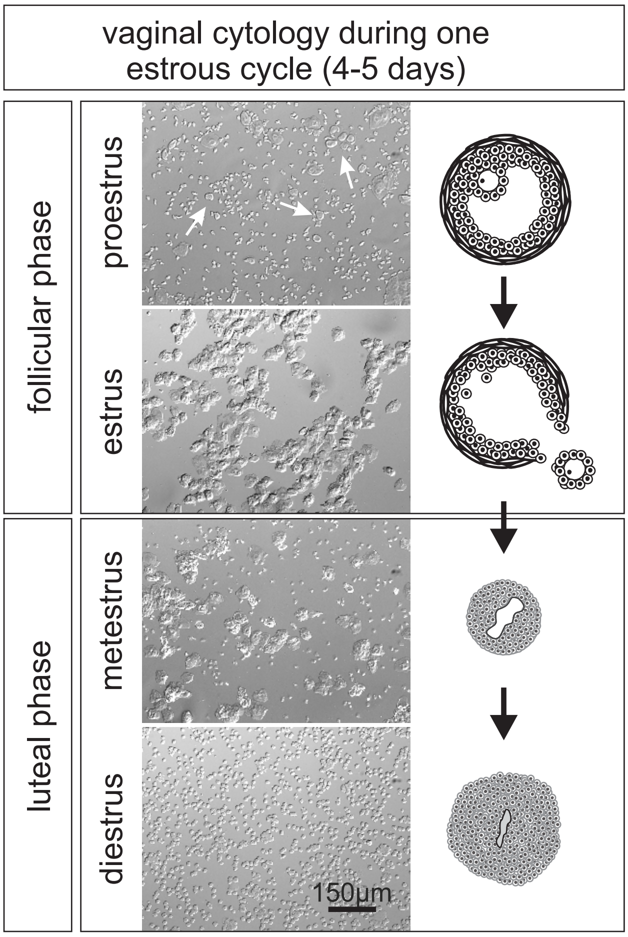


**Supplemental Fig. S1** ***The estrous cycle of mice.*** Smears from vaginal lavage of mice show the distinct cell types of different estrous stages. Proestrus shows multiple nucleated cells (arrows) while the estrus sample only displays cornified epithelial cells. When the mouse enters the luteal phase, leukocytes start to appear and at diestrus they are the dominant cell type in the smear. At proestrus the antral follicle is formed, as depicted in the drawing. The mature oocyte is ovulated at estrus after which the remaining follicle goes through luteinization and forms corpus luteum (CL). Active secretion of progesterone from CL peaks at diestrus followed by luteolysis of the CL cells and resumption of proestrus.

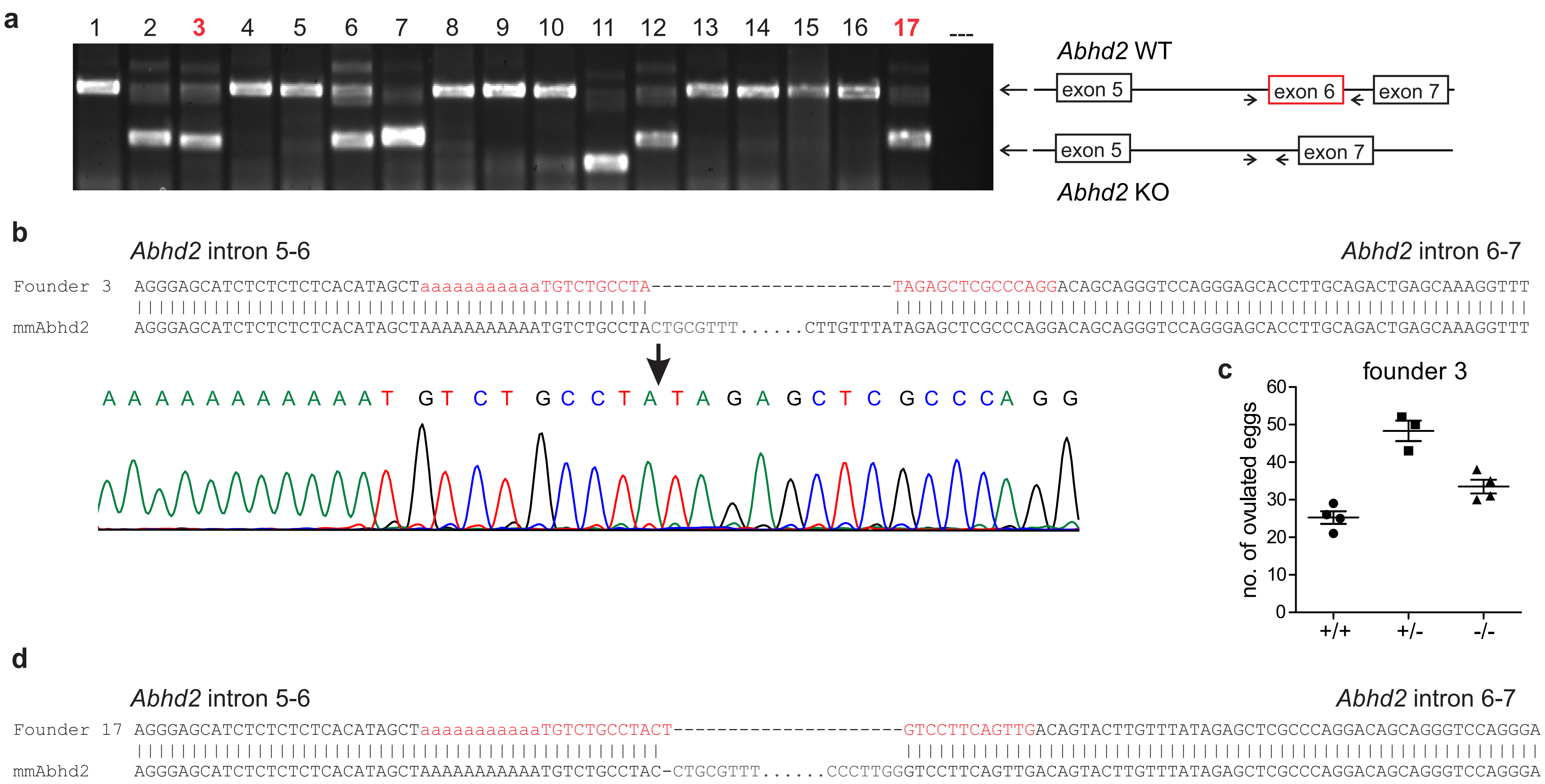

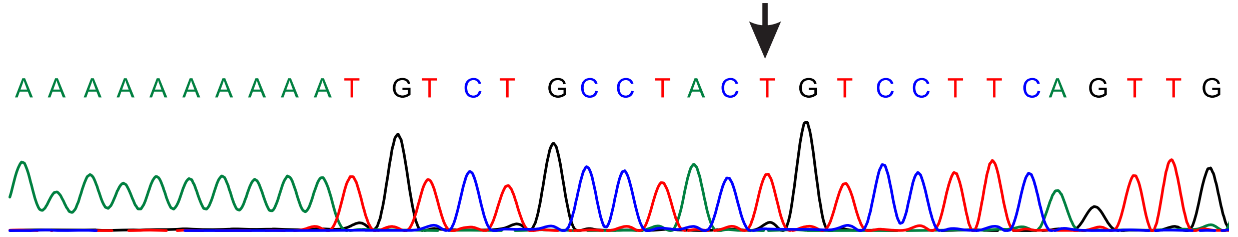

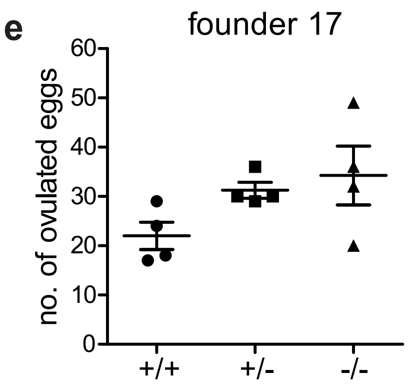


**Supplemental Fig. S2 *CRISPR-EZ results.*** **a)** Genotyping of the 17 mice born after implantation of 78 CRISPR-EZ treated 2-cell stage embryos showed 7 pups with a deletion of *Abhd2* exon 6. Two of the mice, marked in red, were chosen for further phenotypic analysis. **b)** and **d)** Sequencing of the shorter *Abhd2* KO product, obtained after genotyping PCR of the founder mice, showed a deletion of exon 6 starting in the surrounding introns. The chromatograms correspond to the sequence marked in red, with arrows pointing to the ligation site of the two introns. **c)** and **e)** Superovulation results from pups born from the two founder mouse lines display similar increase in number of ovulated eggs.

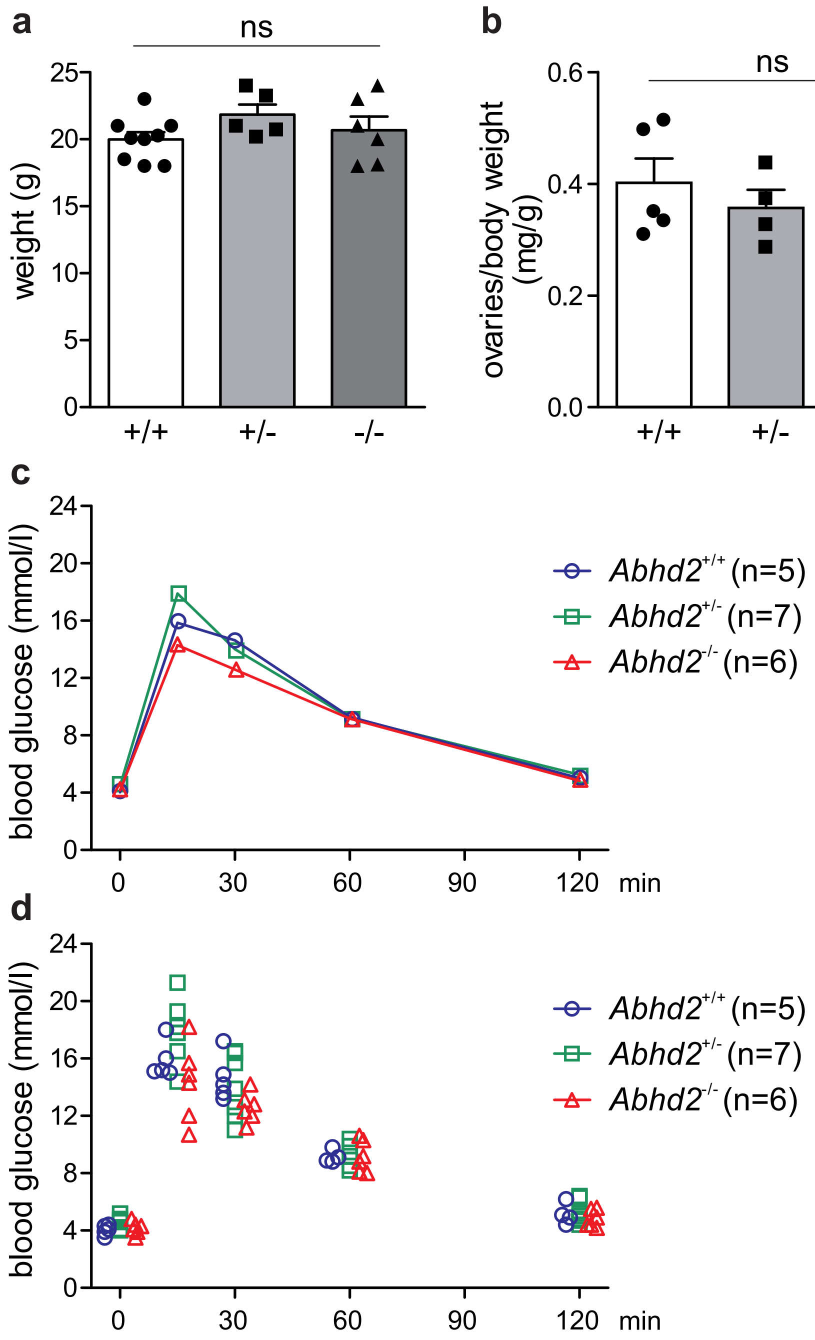


**Supplemental Fig. S3** ***Abhd2 ablation does not alter the weight or the glucose uptake of mice.*** **a)** and **b)** The weight of 2-4-month-old female mice and ovaries from 3-4-month-old mice in proestrus did not differ between genotypes. **c)** Average values of glucose tolerance test of 2.5-month-old female mice, fasted overnight and thereafter given an intraperitoneal injection of 2 g/kg glucose. **d)** The individual measurements of the mice used for the glucose tolerance test in **(c)**. No statistical difference was detected in glucose uptake when comparing *Abhd2*$^{+/+}$, *Abhd2*$^{+/-}$ and *Abhd2*$^{-/-}$ mice.

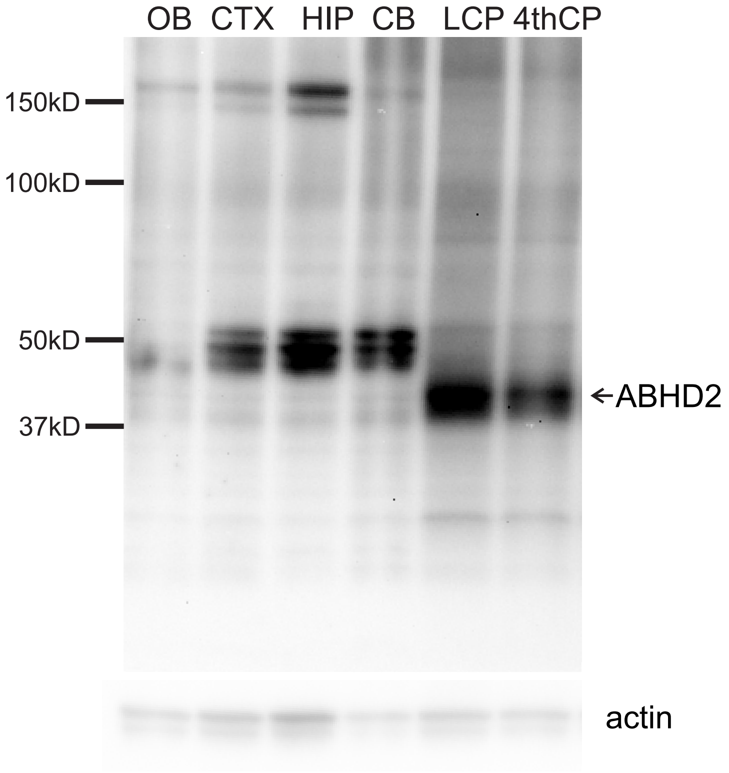


**Supplemental Fig. S4** ***ABHD2 in different brain regions of mice.*** ABHD2 was detected in lateral (LCP) and 4th ventricle choroid plexus (4thCP) by Western blot, while the main olfactory bulb (OB), the cerebral cortex (CTX), the hippocampus (HIP) and the cerebellum (CB) of female mice only show background staining. Samples were collected from a 2.5-month-old *Abhd2*$^{+/+}$ female mouse.
